# Can double PPO mutations exist in the same allele and are such mutants functional?

**DOI:** 10.1101/2022.01.21.477186

**Authors:** Aimone Porri, Matheus M. Noguera, Michael Betz, Daniel Sälinger, Frank Brändle, Steven J. Bowe, Jens Lerchl, Lucie Meyer, Michael Knapp, Nilda Roma-Burgos

## Abstract

**Background:** Resistance to PPO-inhibiting herbicides is primarily endowed by target site mutations at the *PPO2* gene that compromise binding of the herbicide to the catalytic domain. In *Amaranthus* ssp. *PPO2*, the most prevalent target mutations are deletion of the G210 codon and the R128G and G339A substitutions. These mutations strongly affect the dynamic of the PPO2 binding pocket resulting in reduced affinity with the ligand. Here we investigated the likelihood of co-occurrence of the most widespread target site mutations in the same *ppo2* allele.

**Results:** Plants carrying R128G+/+ ΔG210+/-, where + indicates presence of the mutation, were crossed with each other. The *ppo2* of the offspring was subjected to pyrosequencing and *E. coli*- based Sanger sequencing to determine mutation frequencies and allele co-occurrence. The data show that R128G ΔG210 can occur only in one allele; the second allele carries only one mutation. Double mutation in both alleles is less likely because of significant loss of enzyme activity. The segregation of offspring populations derived from a cross between heterozygous plants carrying ΔG210 G399A also showed no co-occurrence in the same allele. The offspring exhibited the expected mutation distribution patterns with few exceptions.

**Conclusions:** Homozygous double-mutants are not physiologically viable. Double-mutant plants can only exist in a heterozygous state. Alternatively, if two mutations are detected in one plant, each mutation would occur in a separate allele.

**Nomenclature:** Palmer amaranth, *Amaranthus palmeri* S. Wats.; protoporphyrinogen oxidase, PPO; tall waterhemp, *Amaranthus tuberculatus* (<u>Moq.</u>) J.D.Sauer

## 1 Introduction

Protoporphyrinogen IX oxidase (PPO, EC. 1.3.3.4) pertains to a highly conserved family of membrane-bound enzymes in the tetrapyrrole biosynthetic pathway that can be found in mammals, plants, bacteria and fungi^1^. It catalyzes the six-atom oxidation of protoporphyrinogen IX to protorphyrin IX, which is the last common step in the production of chlorophylls and heme. In plants, two isoforms encoded by two distinct nuclear genes are found: PPO1 in chloroplasts and PPO2 in the mitochondria.^2^ Some plant species can have PPO2 targeted to both organelles.^3^

PPO-inhibitors have been used as herbicides for more than 60 years. Their high efficacy, broad weed control spectrum, residual activity and the appearance of glyphosate- and ALS-resistant weeds contributed to their large-scale adoption and intensive use by farmers. Eight distinct chemical families of PPO-inhibiting herbicides have been commercially developed so far: diphenyl ethers, triazolinones, thiadiazoles, oxadiazoles, oxazolidinedione, phenylpyrazoles, pyrimidinediones, and N-phenylphthalimides. Much of the variation in herbicide efficacy across PPO-inhibitors can be attributed to their chemical structures, as these molecules bind to PPO by resembling its substrate.

Although the selection of weeds resistant to PPO-inhibitors was a slow process compared to herbicides with other modes of action, biotypes from 13 species have developed resistance mechanisms to these herbicides so far.^4^ The involvement of herbicide metabolism has been reported but not fully characterized,^5, 6^ and a polymorphism in the *ppo1* gene responsible for oxadiazon resistance in *Eleusine indica* was recently discovered.^7^ Nevertheless, mutations in the *ppo2* gene have been identified as the main resistance mechanism to PPO-inhibitors. Among those, the G210 codon-deletion (ΔG210) was the first to be reported, in 2001,^8^ and is so far the most prevalent resistance-conferring mutation among *A. tuberculatus* and *A. palmeri* species.^9-11^ In 2019, a glycine to alanine substitution at the *A. palmeri ppo2* 399^th^ position (G399A) was reported,^12^ and has not been found in other species yet. Interestingly, neither of the native residues (G210 and G399) are directly related to protogen binding to PPO2,^13, 14^ so the resistance caused by mutation at these loci is unexpected. It was later understood that ΔG210 caused an enlargement of the binding pocket compared to the wild-type protein, allowing concomitant binding of inhibitor and substrate.^15^ Protogen binding to ΔG210 PPO2 was not negatively affected, which is a favorable condition to minimize related fitness costs.^15, 16^ However, the same cannot be said about G399A. The authors concluded that the additional methyl group from Ala in relation to Gly protruded into the binding-site, creating repulsive interactions towards the inhibitor (and consequently, towards the substrate).^12^ This mutation led to a 97% reduction in enzyme activity compared to the WT PPO2. Lastly, mutations at the Arg128 position (homologous to Arg98, in *Ambrosia artemisiifolia*) has been reported in several species, such as *A. tuberculatus*,^10^ *A. palmeri*,^17^ *A. retroflexus*,^18, 19^ *A. artemisiifolia*,^20^ and *Euphorbia heterophylla*.^21^ The most common substitution is R128G (4 species), while R128M, R128I and R128L are present in only one species each (*A. palmeri*, *A. tuberculatus* and *A. artemisiifolia*, respectively). The importance of this residue for protogen binding relates to the salt bridge formed between the positively charged Arg with the negatively charged carboxyl group of ring C of protogen.^1, 13, 14, 22^

The continued use of PPO-inhibitors, the dioecious character of some *Amaranthus* species and their hybridization capacity, has led to an accumulation of mutations at the population-, plant- and allele-levels, ^9, 11, 17, 23^ although the latter is very rare. Computational data suggested that both combinations ΔG210 + G399A and G399A + R128G would confer high resistance to fomesafen, but the very low frequency of such genotypes, and the absence of double homozygous plants could also indicate that the resultant protein might be inactive.^9^ In addition, it was predicted that the combination ΔG210 + R128G would not be viable due to the loss of three-dimensional integrity of the binding pocket, which was backed up by the absence of such genotype among the fomesafen-survivors. To assess the herbicide resistance risk arising from double mutations, the aim of this study was to verify which mutation-combinations would be tolerable in PPO2 from *A. palmeri*; and (2) study the inheritance pattern of a double-heterozygous ΔG210 G399A mutant in *A. palmeri*, which was the most prevalent genotype among the double-mutation carriers.

## 2 Materials and Methods

### 2.1. Detection of ppo2 target site mutations by pyrosequencing

The following work was carried out in the laboratory of IDENTXX GmbH (Stuttgart, Germany, www.identxx.com).

*Amaranthus palmeri* plants were sampled by collecting a 0.5 cm² leaf section and individually transferring into collection microtubes (Qiagen, Hilden, Germany). The samples were then homogenized in a shaking mill (TissueLyser II; Qiagen, Hilden, Germany) using steel beads. The DNA extraction was carried out in the KingFisher™ Flex Magnetic Particle Processors (Thermo Fisher Scientific, Schwerte, Germany) using the Chemagic Plant 400 kit (Perkin Elmer, Rodgau, Germany) according to the manufacturer’s instructions (modified by IDENTXX GmbH). The Endpoint PCR amplification (gDNA concentration of 20-50 ng μl^−1^ per sample) for each target region were performed using MyFi™ DNA Polymerase Kit (Bioline GmbH, Luckenwalde, Germany) and specific primers (IDENTXX GmbH)in a PCR thermal cycler (T100 PCR thermal cycler, Bio-Rad Laboratories GmbH, Germany) under the following conditions: 3 min at 95°C and 42 cycles of 10 s denaturation at 95°C; 35 s annealing at 60°C and 45 s elongation at 72°C; and a final elongation step at 72°C for 5 min.. The successful amplification (225 bp for the ΔG210, 250 bp for the G399A product) was checked per gel electrophoresis on a 1.5 % agarose gel.

The PCR products were analyzed for SNPs at the target positions via pyrosequencing on a PyroMark Q24 (Qiagen, Hilden, Germany) using specific sequencing primers (IDENTXX GmbH). During the sequencing reaction, all incorporated nucleotides of a short region that encompasses the SNP position of interest were detected and reported by creating a pyrogramm in a pyrorun file. The created file was read by the PyroMark Q24 software (Version 2.0.7) and visually evaluated for mutations, both heterozygous and homozygous.

### 2.2. Verification of double mutants by cloning in *E. coli*

The following work was carried out in the laboratory of IDENTXX GmbH (Stuttgart, Germany, www.identxx.com).

A 0.5 cm² of leaf sample of each plant (continuously cooled at −20°C) was transferred into a collection microtubes. The samples were then homogenized at room temperature in a shaking mill. The RNA extraction was carried out in the KingFisher™ Flex Magnetic Particle Processors (Thermo Fisher Scientific, Schwerte, Germany) using the MagMAX™ Plant RNA Isolation Kit (Thermo Fisher Scientific Inc, Schwerte, Germany) according to the manufacturer’s instructions (modified by IDENTXX GmbH). The cDNA transcription was done using the High-Capacity cDNA Reverse Transcription Kit (Thermo Fisher Scientific Inc, Schwerte, Germany) according to the manufacturer’s instructions (modified by IDENTXX GmbH). The Endpoint PCR amplification for the target region encompassing both SNPs positions were performed using MyFi™ DNA Polymerase Kit and specific primers (IDENTXX GmbH) in a PCR thermal cycler under the following conditions: 5 min at 95°C and 42 cycles of 20 s denaturation at 95°C; 35 s annealing at 60°C and 90 s elongation at 72°C; and a final elongation step at 72°C for 5 min. The successful amplification of the 1850 bp PCR product was checked per gel electrophoresis on a 1.5 % agarose gel.

PCR products were cloned using StrataClone PCR Cloning Kit (Agilent, Waldbronn, Germany). Positive white colonies were randomly picked and verified with colony PCR. For each clone, 10 positive PCR fragments were randomly selected and verified via Sanger sequencing (SeqLab-Microsynth, Göttingen, Germany). Sequences were analyzed using Geneious software v. 9.1.8 (Biomatters, Auckland, New Zealand).

### 2.3. Evaluation of mutant PPO enzyme activity

The complete description of expression and purification of Amaranthus PPO2 variant proteins and the enzymatic assay to determine protein activity was referenced to the method described by Rangani et al.^12^ To calculate the percentage of remaining protein activity, the enzyme activity of WT PPO2 was divided by the activities of R128G, ΔG210 and R128G ΔG210 PPO2 variants and multiplied by 100. This was further normalized to the amount (ng) of protein used in the assay.

### 2.4 Arabidopsis transgenics growth and herbicide treatment

#### 2.4.1 Donor plant material and growth conditions

*Arabidopsis thaliana* seeds (stock MC24, from the Max Planck Institute for Molecular Plant Physiology at Golm) were sown into a substrate composed of GS90 soil + 5% sand. Plants were subjected to stratification for 5 days at 4°C, followed by a short-day growth period of 10 d (10/14 h of day/night intervals at 20/18°C ± 1°C, ~120 μmol PAR). After that, plants were transplanted into 8X8 cm pots filled with GS90 soil and cultivated under the same conditions for 14 d, under long-day growth conditions (16/8 h day/night at 20/18°C ± 1°C, ~200 μmol PAR) and maintained until seed harvest. Plants were fertilized with 0.3% Hakaphos Blau (15-10-15 NPK) twice a week until flowering. Relative humidity was not controlled, but kept between 40-70% during all growth stages, except during stratification.

#### 2.4.2 Transgene preparation

To prepare the transgene, the mutant *ppo2* was inserted into RTP6557 transformation vector, which was then inserted into *Agrobacterium tumefaciens* strain C58C1pMP90. The gene insert also included an acetolactate synthase-herbicide-resistance trait as a selectable marker to identify transformed *Arabidopsis* seedlings. This would ensure that plants eventually tested for resistant to PPO herbicide all expressed the transgene.

#### 2.4.3 Bacterial culture and dipping medium

*Agrobacterium* culture containing the plasmid was prepared a day before dipping by inoculating 1 ml of glycerol stock into 250 ml of YEB medium (1 g L^−1^ yeast, 5 g L^−1^ beef extract, 5 g L^−1^ peptone, 5 g L^−1^ sucrose, 0.49 g L^−1^ MgSO_4_*7H_2_O) + appropriate antibiotic. The bacteria were cultured for 12 h at 28°C with continuous agitation at 150 rpm. The next day, after adjusting the *Agrobacterium* culture density to OD600 = 1.0 (with YEB medium), the culture was collected by centrifugation at 4000 rpm for 10 min and re-suspended in 150 ml *infiltration medium* composed of 2.2 g L^−1^ MS (Murashige & Skoog medium), 50 g L^−1^ sucrose, 0.5 g L^−1^ MES hydrate, 10μl L^−1^ BAP (Benzylaminopurin, 1 mg ml^−1^). The pH was then adjusted to 5.7-5.8.

#### 2.4.4 Agrobacterium-mediated transformation of Arabidopsis thaliana by floral dip

Plant transformation was performed following a previously established protocol. ^24^ Briefly, plants with immature floral buds were dipped in the bacterial suspension for 10 sec after adding 75 μl of Silwet-L77 per 150 ml of infiltration medium to a jar. After dipping, plants were kept overnight in a cabinet under high humidity and low light intensity, and were grown under long-day conditions until maturity. When siliques turned yellow, plants were placed inside paper bags to collect the seeds. T1 seeds were then transferred to falcon tubes and stored at 4°C.

#### 2.4.5 Selection of putative transformants with Imazamox

After 14 or more days of storage at 4°C, T1 seeds were sown to select putative transgenic *Arabidopsis* plants. Sowing and stratification was performed as described previously. Following that, seeds were treated with a 20 ppm imazamox solution and cultivated under short-day growth conditions for 12-14 days, when resistant seedlings (4-leaf stage) were transplanted into 6 cm pots filled with GS90 soil and grown for another 10 days. One day prior to herbicide application, growth conditions were set to ‘long-day’ and maintained throughout the duration of the test. Herbicide treatments consisted of two concentrations of saflufenacil (Kixor, BASF Corporation, at 10 and 25 g ai ha^−1^) foliar-applied when plants reached the 10-leaf stage, using a spray chamber calibrated to deliver 375 L ha^−1^ of spray solution. Herbicide efficacy was visually assessed after 7 days from herbicide treatments.

### 2.5. Verifying PPO double mutants from heterozygous dG210 x heterozygous G399A

In a previous study we learned that ΔG210 −/+ G399A −/+ was the most common genotype among the double mutant plants identified.^9^ The double mutants referred to here were plants carrying two *ppo2* mutations, but without confirmation on whether the two mutations occurred in one allele. To study the inheritance pattern of this genotype, seeds from the selected Palmer amaranth population were sown in a plastic tray containing potting mix (Sunshine LC1; SunGro Horticulture, Agawam, MA, USA). After one week, 400 seedlings were transplanted to 50-cell trays at one seedling per cell. When plants reached the 6-to 8-cm stage, fomesafen (264 g ai ha^−1^, Flexstar® 1.88 EC, Syngenta Crop Protection, Greensboro, NC, USA) was applied in a spray chamber, equipped with two flat-fan 100 0065 nozzles, calibrated to deliver 187 L ha^−1^ of spray mix at a speed of 3.6 km h^−1^. At 21 d after application, leaf tissues from around 200 survivors were individually collected into 1.5-mL microtubes (VWR International LLC, Radnor, PA, USA) and stored at −80 ◻ until processing. DNA was extracted and the plants were genotyped via pyrosequencing as previously described. Four males and 14 females of the same genotype (ΔG210 −/+ G399A −/+) were individually transplanted to 8-L pots and grown together to interbreed. Seeds were harvested separately from each female plant and cleaned. The germination capacity of 14 F1 lines was evaluated. Six F1 lines with the higher germination capacity were selected, and up to 100 plants from each female were selected for genotyping via pyrosequencing.

Since Palmer amaranth is diploid, a plant that is heterozygous for both mutations can either: a) have one allele containing two mutations, while the other allele is WT; or b) have each allele carrying one mutation only. The Palmer amaranth parent population used in this study exhibited the latter condition. A cross between two double-heterozygous is expected to result in 25% of the offspring carrying a homozygous ΔG210, 25% carrying a homozygous G399A and 50% of the offspring having the same parental genotype. A Pearson’s chi-squared test was used to confirm this hypothesis, where the calculated chi-squared value (χ^2^ Calc, as shown in Equation 1) is compared to a tabulated value (χ^2^ Tab)

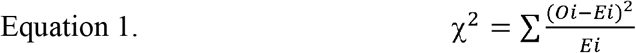

where *Oi* = observed number of plants of a genotype and *Ei* = expected number of plants of a genotype.

Lastly, the occurrence of double mutants was verified by cloning the *ppo2* gene into *E. coli* as previously described. A total of 26 samples were chosen for this procedure, and around 10 clones from each sample were Sanger-sequenced.

## 3 Results and Discussion

### 3.1. PPO mutation profile of R128G+/+ ΔG210-/+ offspring

A male and a female plant LIH11640A and LIH11640F carrying R128G+/+ ΔG210+/− (where + indicates presence of the mutation), were detected in a large biotype screening of putative resistant *A. palmeri*. The R128G+/+ ΔG210+/− plants carried homozygous R128G, indicating that double mutations ΔG210 and R128G can occur on the same allele. This finding answers the question we posed in a previous study,^9^ where this genotype was not observed among the double mutants. In that study, we hypothesized that: a) either the occurrence of this genotype was too rare and would require a bigger sample size to be detected; or b) the resultant protein would be inactive and, therefore, the fitness cost associated with this double-mutation would be lethal.

To determine whether homozygous double mutants R128G ΔG210 plants are viable, LIH11640A was crossed with LIH11640F and the *ppo2* of LIH11640F1 offspring was subjected to pyrosequencing for the positions R128 and G210. If the R128G ΔG210 PPO is functional, and inheritance follows a Mendelian pattern, it is expected that two parents having the R128G+/+ ΔG210+/− genotype would result in 25% of the offspring being double-homozygous, 25% being R128G+/+ G210 and 50% having the parental genotype (Fig 1).

**Figure 1.**
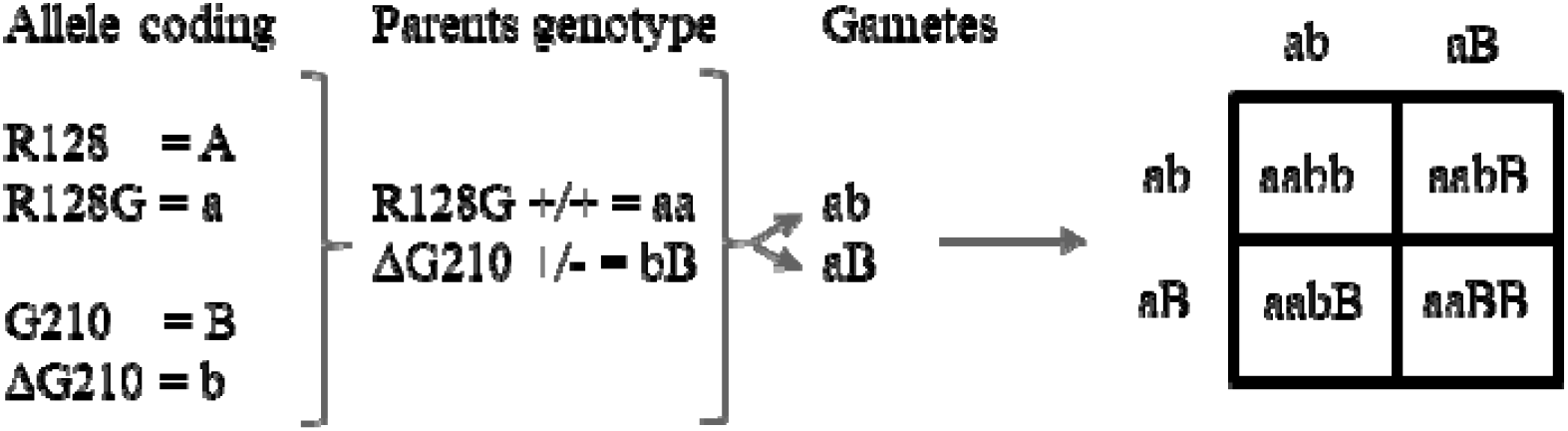
The expected segregation pattern of the offspring from a cross of homozygous R128G mutant (R128G+/+) and heterozygous ΔG210 mutant (ΔG210+/−).

Out of 100 plants genotyped from the LIH11640F1 population, only 29% of the offspring carried the parental genotype; 71% of the plants were R128G +/+ G210; and none were homozygous double mutants (Table 1). The calculated chi-squared value for this population was three times higher than the tabulated (χ^2^ Calc = 18.46 and χ^2^ Tab = 5.991), indicating that observed and expected ratios do not agree. Due to the odd inheritance pattern in LIH11640F1, additional R128G+/+ ΔG210+/− male and female plants from this population were selected and crossed to generate another three independent LIH11640F2 populations, which were also pyrosequenced (Table 1). The mutation inheritance pattern among the three F2 populations conformed to that of the F1 population, where no homozygous double mutants were detected and the majority of the offspring carried a single, homozygous R128G mutation (R128G +/+ G210). All calculated chi-squared values were higher than the tabulated. These results indicate that, although rare individuals containing both R128G ΔG210 mutations can occur, their co-occurrence on both alleles is not possible. In addition, the percentage of F1 and F2 plants carrying R128G+/+ ΔG210+/− was significantly lower than the expected ratio (Table 1), suggesting that having R218G and ΔG210 in one allele has a significant impact on fitness cost.

**Table 1.**
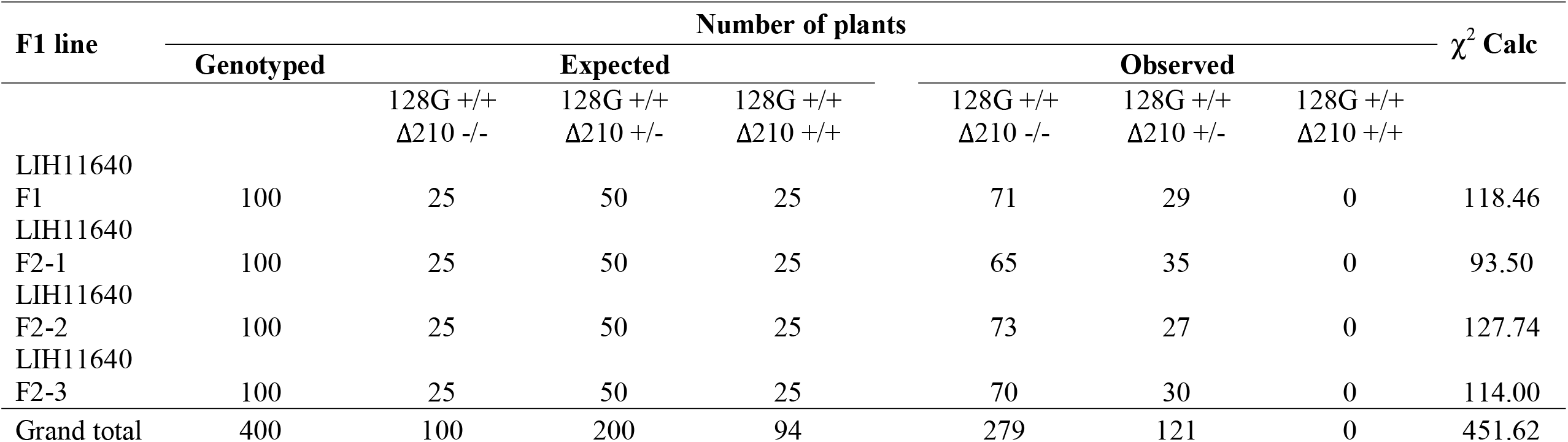
Segregation of mutant alleles in F1 and F2 generations of PPO-herbicide-resistant *Amaranthus palmeri* genotypes.

### 3.2. Verification of double mutant allele combination by cloning in E. coli and Sanger sequencing

For further verification of the pyrosequencing results, the plants coded as LIH11640A and LIH11640F were used for RNA extraction, cDNA synthesis, and amplification of the whole-length *PPO2* gene, which was later cloned into *E. coli*. For each plant, 10 clones were selected for Sanger sequencing.

As expected, all clones from both plants carried the R128G mutation. The number of clones containing double mutations was similar for both plants studied: four and five out of 10 clones for plants LIH11640A and LIH11640F, respectively (Table 2). Since the *ppo2* fragments inserted into *E. coli* were derived from RNA of the parent plants, the balanced distribution of double- and single-mutants suggests that there is no difference in the expression of those alleles. In other words, LIH11640A and LIH11640F plants do not favor the expression of one allele over the other, and the amount of double- and single-mutant proteins produced *in vivo* are equivalent.

**Table 2.** Confirmation of the occurrence of R128G and ΔG210 mutants in PPO2 clones in *E. coli* from PPO-herbicide-resistant *Amaranthus palmeri* plants.

| Plant ID | Clone ID | R128G | ΔG210 |
| --- | --- | --- | --- |
| AMAPA LIH11640A | Clone 1 | + | + |
|  | Clone 2 | + | - |
|  | Clone 3 | + | + |
|  | Clone 4 | + | - |
|  | Clone 5 | + | - |
|  | Clone 6 | + | - |
|  | Clone 7 | + | + |
|  | Clone 8 | + | - |
|  | Clone 9 | + | - |
|  | Clone 10 | + | + |
| AMAPA LIH11640F | Clone 1 | + | + |
|  | Clone 2 | + | - |
|  | Clone 3 | + | - |
|  | Clone 4 | + | + |
|  | Clone 5 | + | - |
|  | Clone 6 | + | - |
|  | Clone 7 | + | + |
|  | Clone 8 | + | - |
|  | Clone 9 | + | + |
|  | Clone 10 | + | + |

### 3.3. Evaluation of double-mutant R128G ΔG210 PPO2 enzymatic activity

The segregation of mutations in the offspring suggests that the double mutant (R128G ΔG210) protein activity is either severely compromised or non-existent, and plants carrying such double-mutant allele rely exclusively on the other, single-mutant allele to survive. To confirm this hypothesis, the WT, single-mutant and double-mutant enzymes were heterologously expressed in *E. coli*, the PPO2 protein was purified, and the enzyme activity was quantified. Enzyme activity data are shown as percentages relative to the WT protein (Table 3). Of the three mutants tested, the R128G variant showed the highest remaining activity, followed by the ΔG210 and the double-mutant variants. The enzyme activity of the double mutant was well below the detection limit, indicating that it was practically inactive. The inactivity of the double-mutant enzyme agrees with our previous study,^9^ where we deduced that a R128G ΔG210 protein would lose its binding pocket integrity.

**Table 3.** Enzyme activity assay of *Amaranthus palmeri* PPO2 mutants.

| **PPO Variant** | **FU/min** | **% Remaning activity** |
| --- | --- | --- |
| WT PPO2 (25ng) | 584 | 100 |
| ΔG210 (500ng) | 1500 | 11 |
| R128G (25ng) | 250 | 50 |
| R128G ΔG210 (500 ng) | 15 | 0.12 |

The effect of ΔG210 and R128G mutations on PPO activity has already been investigated and our results somewhat agrees with previous studies.^15, 16, 18, 25^ These mutations are known to affect PPO activity and substrate binding differently. While ΔG210 did not affect substrate binding (no differences in Km between WT and mutant), the mutation reduced enzyme activity by a factor of 10.^15^ The deletion of G210 causes an enlargement of the substrate binding pocket and displaces G207, which has an important role in positioning protogen at the ideal distance and orientation from the co-factor FAD, favoring the rate-limiting hydride abstraction.^15^ The loss of that interaction results in lower protein activity. In contrast, R128G impairs protein binding efficacy, but doubles its activity.^18^ The decrease in substrate binding is caused by the loss of the salt bridge interaction between R128 and protogen. However, this also facilitates the release of proto from the binding site. In addition, the substitution of the bulky, charged Arg to the small, non-polar Gly, increases the opening of the binding pocket, further speeding the proto egress.^18,25^

To confirm the double-mutant viability, *Amaranthus palmeri* PPO2 carrying R128G ΔG210 was overexpressed in *Arabidopsis* and the resulting transgenic plants were treated with two rates of saflufenacil (10 and 25 g ha^−1^). If active, transgenic plants expressing R128G ΔG210 should be resistant to saflufenacil. The R128G ΔG210 *Arabidopsis* transgenics did not survive the field dose of saflufenacil, indicating that protein activity of this variant is too low to confer herbicide tolerance (Fig 2). The absence of a significant tolerance of the transgenic line suggests that R128G ΔG210 strongly impairs PPO2 protein function.

**Figure 2.**
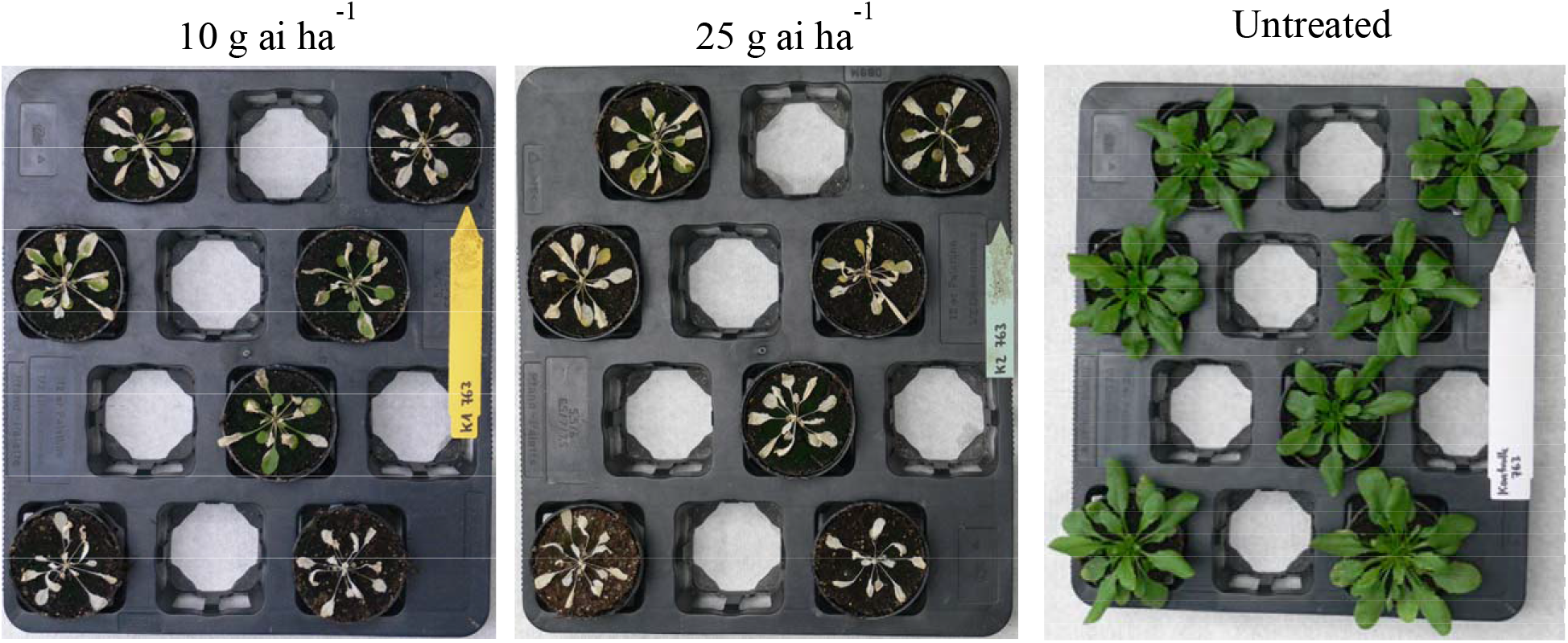
T1 35S:AMAPA PPO2 R128G ΔG210 Arabidopsis lines sprayed with Saflufenacil. Pictures were taken 7 days after treatment. The bottom 2 pots contain wild-type plants, the upper 5 pots contain independent transgenic events (T1 plants, selected with Imazamox by confirming presence of resistance gene AHAS).

### 3.4. PPO mutation profile of crosses between double heterozygous ΔG210G399A

Among field populations of Palmer amaranth, the most common *ppo2* mutation is ΔG210 and the most common mutations detected in the same population are ΔG210 and G399A.^9^ Therefore, it is only logical to assume that due to the obligate outcrossing behavior of this specie, the co-existence of single-mutant plants (regardless of zygosity) in one population would only increase the frequency of ΔG210 G399A double mutants with time. This could then become the most common resistant genotype that would challenge current and future PPO-inhibiting herbicides. That is, if such double mutant does not carry fitness penalty. Hence this follow-up experiment. We expected that a cross between two double-heterozygous ΔG210 G399A would result in 50% of the offspring carrying a single homozygous mutation (either ΔG210 or G399A), while the other 50% would carry both mutations concomitantly as heterozygous. To check if the observed values agreed with the expected inheritance pattern, a chi-squared test was done. The χ^2^ Calc values for individual F1 lines ranged from 1.02 to 4.73, and were lower than the tabulated threshold (χ^2^ Tab = 5.991), confirming that the observed inheritance pattern agrees with the expected (Table 4). The same result was obtained when considering samples across all F1 lines (N=376), where χ^2^ Calc = 2.33.

**Table 4:** Genotyping of Palmer amaranth *ppo2* mutations via pyrosequencing from six F1 lines produced by a cross between double-heterozygous ΔG210/G399A parents

| F1 line | Number of plants | | | | | | | $\chi^2$ Calc |
| --- | --- | --- | --- | --- | --- | --- | --- | --- |
|  | Genotyped | Expected |  |  | Observed |  |  |  |
| | | ΔG210 -/-<br>G399A +/+ | ΔG210 +/-<br>G399A +/- | ΔG210 +/+<br>G399A -/- | ΔG210 -/-<br>G399A +/+ | ΔG210 +/-<br>G399A +/- | ΔG210 +/+<br>G399A -/- | |
| LIH19623 | 52 | 13 | 26 | 13 | 14 | 19 | 19 | 4.73 |
| LIH19624 | 93 | 23.3 | 46.5 | 23.3 | 16 | 51 | 26 | 3.02 |
| LIH19625 | 97 | 24.3 | 48.5 | 24.3 | 21 | 56 | 20 | 2.34 |
| LIH19626 | 20 | 5 | 10 | 5 | 3 | 10 | 7 | 1.60 |
| LIH19627 | 14 | 3.5 | 7.0 | 3.5 | 3 | 10 | 1 | 3.14 |
| LIH19628 | 100 | 25 | 50 | 25 | 28 | 51 | 21 | 1.02 |
| Grand total | 376 | 94 | 188 | 94 | 85 | 197 | 94 | 2.33 |

A total of 26 plants double-heterozygous from all the F1 lines were selected for cloning of the *PPO2* gene into *E. coli*. At least 10 clones per plant were submitted for Sanger sequencing. In this case, half of the clones were expected to contain the G210 deletion, while the other half would carry G399A. This distribution was tested using the Pearson’s chi-squared goodness-of-fit test. Four out of six F1 lines had the observed distribution equal to the expected, but the lines LIH19626 and LIH19628 did not fit this pattern (Table 5). In these two cases, the higher number of clones containing the ΔG210 mutation indicates an unequal expression of this allele in comparison to the allele containing G399A. The mechanism or reason for this phenomenon is not yet understood.

**Table 5:** Sanger sequencing of *E. coli* clones expressing the *ppo2* gene from six F1 *Amaranthus palmeri* lines

| F1 line | # of plants<br>cloned | Number of clones | | | | | $\chi^2$ Calc |
| --- | --- | --- | --- | --- | --- | --- | --- |
|  |  | Sequenced | Expected |  | Observed |  |  |
|  |  |  | dG210 | G399A | dG210 | G399A |  |
| LIH19623 | 6 | 62 | 31 | 31 | 30 | 28 | 0.067 |
| LIH19624 | 3 | 30 | 15 | 15 | 17 | 17 | 0.267 |
| LIH19625 | 3 | 30 | 15 | 15 | 17 | 13 | 0.267 |
| LIH19626 | 5 | 51 | 25.5 | 25.5 | 35 | 16 | 7.078 |
| LIH19627 | 3 | 36 | 18 | 18 | 15 | 21 | 1.000 |
| LIH19628 | 6 | 62 | 31 | 31 | 40 | 22 | 5.225 |

## 4 CONCLUSIONS

The co-occurrence of two mutations in a same *ppo2* allele, although rare, is possible. However, the coded protein has a severely reduced activity, which is supported by the fact that no double-homozygous plants were detected, and that its insertion in Arabidopsis did not result in resistance to PPO-inhibitors. Plants carrying a double-mutant allele must rely on the alternative allele to produce a functional enzyme. The fitness cost in plants carrying a double-mutant allele is yet to be determined.

## Supporting information

Graphical abstract text

## 5 ACKNOWLEDGEMENTS

We are grateful to Daniel Sälinger for providing the PPO enzyme substrate and we thank Hardy Schön and Sarah Heyn for the Arabidopsis service transformation.

## 6 CONFLICT OF INTERESTS

Authors affiliated with BASF have contributed to the planning and implementation of research activities. All other authors declare no conflict of interest.

---

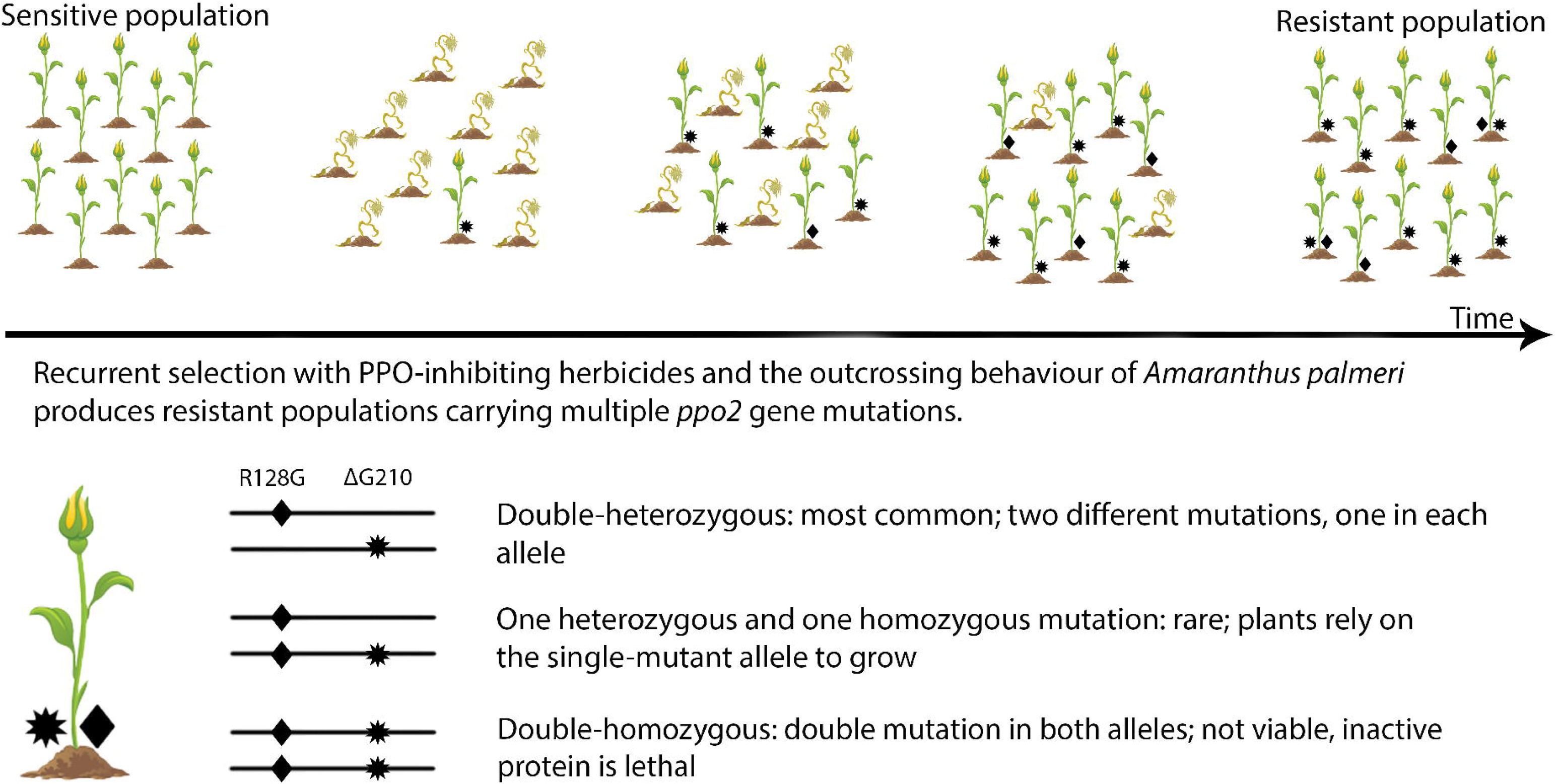

## Notes

### Competing Interest Statement

The authors have declared no competing interest.

## REFERENCES

1. Hao G-F, Zuo Y, Yang S-G and Yang G-F, Protoporphyrinogen oxidase inhibitor: an ideal target for herbicide discovery. CHIMIA International Journal for Chemistry 65: 961-969 (2011).

2. Lermontova I, Kruse E, Mock H-P and Grimm B, Cloning and characterization of a plastidal and a mitochondrial isoform of tobacco protoporphyrinogen IX⍰oxidase. Proceedings of the National Academy of Sciences 94: 8895-8900 (1997).

3. Watanabe N, Che F-S, Iwano M, Takayama S, Yoshida S and Isogai A, Dual Targeting of Spinach Protoporphyrinogen Oxidase II to Mitochondria and Chloroplasts by Alternative Use of Two Inframe Initiation Codons *. J Biol Chem 276: 20474-20481 (2001).

4. Heap I, International Survey of Herbicide Resistant Weeds. www.weedscience.org [accessed Jan 7 2021].

5. Obenland OA, Ma R, O’Brien SR, Lygin AV and Riechers DE, Carfentrazone-ethyl resistance in an Amaranthus tuberculatus population is not mediated by amino acid alterations in the PPO2 protein. PloS one 14: e0215431 (2019).

6. Varanasi VK, Brabham C and Norsworthy JK, Confirmation and Characterization of Non–target site Resistance to Fomesafen in Palmer amaranth (Amaranthus palmeri). Weed Sci 66: 702-709 (2018).

7. Bi B, Wang Q, Coleman JJ, Porri A, Peppers JM, Patel JD, et al., A Novel Mutation A212T in Chloroplast Protoporphyrinogen Oxidase (PPO1) Confers Resistance to PPO Inhibitor Oxadiazon in Eleusine indica. Pest Manage Sci 2019).

8. Patzoldt WL, Hager AG, McCormick JS and Tranel PJ, A codon deletion confers resistance to herbicides inhibiting protoporphyrinogen oxidase. Proc Natl Acad Sci USA 103: 12329-12334 (2006).

9. Noguera MM, Rangani G, Heiser J, Bararpour T, Steckel LE, Betz M, et al., Functional PPO2 mutations: co-occurrence in one plant or the same ppo2 allele of herbicide-resistant Amaranthus palmeri in the US mid-south. Pest Manage Sci 2020).

10. Nie H, Mansfield BC, Harre NT, Young JM, Steppig NR and Young BG, Investigating target-site resistance mechanism to the PPO-inhibiting herbicide fomesafen in waterhemp and interspecific hybridization of Amaranthus species using next generation sequencing. Pest Manage Sci 75: 3235-3244 (2019).

11. Copeland JD, Giacomini DA, Tranel PJ, Montgomery GB and Steckel LE, Distribution of PPX2 Mutations Conferring PPO-Inhibitor Resistance in Palmer Amaranth Populations of Tennessee. Weed Technol 32: 592-596 (2018).

12. Rangani G, Salas-Perez RA, Aponte RA, Knapp M, Craig IR, Mietzner T, et al., A Novel Single-Site Mutation in the Catalytic Domain of Protoporphyrinogen Oxidase IX (PPO) Confers Resistance to PPO-Inhibiting Herbicides. Fron Plan Sci 10 2019).

13. Heinemann Ilka U, Diekmann N, Masoumi A, Koch M, Messerschmidt A, Jahn M, et al., Functional definition of the tobacco protoporphyrinogen IX oxidase substrate-binding site. Biochem J 402: 575-580 (2007).

14. Koch M, Breithaupt C, Kiefersauer R, Freigang J, Huber R and Messerschmidt A, Crystal structure of protoporphyrinogen IX oxidase: a key enzyme in haem and chlorophyll biosynthesis. The EMBO Journal 23: 1720-1728 (2004).

15. Dayan FE, Daga PR, Duke SO, Lee RM, Tranel PJ and Doerksen RJ, Biochemical and structural consequences of a glycine deletion in the α-8 helix of protoporphyrinogen oxidase. Biochim Biophys Acta 1804: 1548-1556 (2010).

16. Wu C, Goldsmith M-R, Pawlak J, Feng P, Smith S, Navarro S, et al., Differences in efficacy, resistance mechanism and target protein interaction between two PPO inhibitors in Palmer amaranth (Amaranthus palmeri). Weed Sci 68: 105-115 (2020).

17. Giacomini DA, Umphres AM, Nie H, Mueller TC, Steckel LE, Young BG, et al., Two new PPX2 mutations associated with resistance to PPO-inhibiting herbicides in Amaranthus palmeri. Pest Manage Sci 73: 1559-1563 (2017).

18. Du L, Li X, Jiang X, Ju Q, Guo W, Li L, et al., Target-Site Basis for Fomesafen Resistance in Redroot Pigweed (Amaranthus retroflexus) from China. Weed Sci 69: 290-299 (2021).

19. Huang Z, Cui H, Wang C, Wu T, Zhang C, Huang H, et al., Investigation of resistance mechanism to fomesafen in Amaranthus retroflexus L. Pestic Biochem Physiol 165: 104560 (2020).

20. Rousonelos SL, Lee RM, Moreira MS, VanGessel MJ and Tranel PJ, Characterization of a common ragweed (Ambrosia artemisiifolia) population resistant to ALS-and PPO-inhibiting herbicides. Weed Sci 60: 335-344 (2012).

21. Mendes RR, Takano HK, Adegas FS, Oliveira RS, Gaines TA and Dayan FE, Arg-128-Leu target-site mutation in PPO2 evolves in wild poinsettia (Euphorbia heterophylla) with cross-resistance to PPO-inhibiting herbicides. Weed Sci 68: 437-444 (2020).

22. Hao G-F, Tan Y, Xu W-F, Cao R-J, Xi Z and Yang G-F, Understanding Resistance Mechanism of Protoporphyrinogen Oxidase-Inhibiting Herbicides: Insights from Computational Mutation Scanning and Site-Directed Mutagenesis. J Agric Food Chem 62: 7209-7215 (2014).

23. Lillie KJ, Giacomini DA, Green JD and Tranel PJ, Coevolution of resistance to PPO inhibitors in waterhemp (Amaranthus tuberculatus) and Palmer amaranth (Amaranthus palmeri). Weed Sci 67: 521-526 (2019).

24. Clough SJ and Bent AF, Floral dip: a simplified method for Agrobacterium -mediated transformation of Arabidopsis thaliana. The Plant Journal 16: 735-743 (1998).

25. Hao G-F, Tan Y, Yang S-G, Wang Z-F, Zhan C-G, Xi Z, et al., Computational and Experimental Insights into the Mechanism of Substrate Recognition and Feedback Inhibition of Protoporphyrinogen Oxidase. PLOS ONE 8: e69198 (2013).

